# Identification of novel mutations associated with cycloserine resistance in *Mycobacterium tuberculosis*

**DOI:** 10.1101/114116

**Authors:** Shuo Zhang, Jiazhen Chen, Peng Cui, Wanliang Shi, Wenhong Zhang, Ying Zhnag

**Author notes:** Corresponding author: Jiazhen Chen, PhD, Ying Zhang, MD, PhD. These authors contributed equally to this work (Shuo Zhang, Jiazhen Chen).

## Abstract

**Objectives:** D-cycloserine (DCS) is an important second-line drug used to treat multi-drug resistant (MDR) and extensively drug-resistant (XDR) tuberculosis. However, the mechanisms of resistance to DCS are not well understood. Here we investigated the molecular basis of DCS resistance using in vitro isolated resistant mutants of *Mycobacterium tuberculosis.*

**Methods:** *M. tuberculosis* H37Rv was subjected to mutant selection on 7H11 agar plates containing varying concentrations of DCS. A total of 35 DCS-resistant mutants were isolated and 18 mutants were subjected to whole genome sequencing. The identified mutations associated with DCS resistance were confirmed by PCR-Sanger sequencing.

**Results:** We identified mutations in 17 genes that are associated with DCS resistance. Except mutations in *alr* (*rv3423c*) which is known to be involved in DCS resistance, 16 new genes *rv0059, betP (rv0917), rv0221, rv1403c, rv1683, rv1726, gabD2 (rv1731), rv2749, sugI* (*rv3331*)*, hisC2* (*rv3 772*), single mutation in 5’ intergenic region of *rv3345c* and *rv1435c,* and insertion in 3’ region of *rv0759c* were identified as solo mutations in their respective DCS-resistant mutants. Our findings indicate that the mechanisms of DCS resistance are more complex than previously thought and involve genes participating in different cellular functions such as lipid metabolism, methyltransferase, stress response, and transport proteins.

**Conclusions:** New mutations in diverse genes associated with DCS are identified, which shed new light on the mechanisms of action and resistance of DCS. Future studies are needed to verify these findings in clinical strains so that molecular detection of DCS resistance for improved treatment of MDR-TB can be developed.

## Introduction

D-Cycloserine (DCS) is a cyclic analog of D-alanine and is a broad-spectrum antibiotic that inhibits the growth of Gram-positive and Gram-negative bacteria. Although DCS has psychiatric and nervous system adverse reactions, it displays no cross-resistance with any other known antitubercular drugs ^1^ DCS is an important second-line drug for the treatment of multi-drug resistant (MDR) and extensively drug-resistant (XDR) tuberculosis ^1^, and is currently classified as a Group C agent for the treatment of MDR-TB treatment by WHO ^2^.

Since DCS is a structural analog of D-alanine, enzymes whose substrates are D-alanine are the drug targets in mycobacteria ^3–5^. These enzymes include D-alanine racemase (Alr) and d-alanine:d-alanine ligase (Ddl), which are required for the synthesis of peptidoglycan in the mycobacterial cell wall. Overexpression of *alr* and *ddl* has been shown to cause resistance to DCS in *M. smegmatis* ^6,7^ Moreover, single nucleotide polymorphisms in these genes were also found in resistant *M. tuberculosis* ^8,10^. Consistent with the cell wall peptidoglycan being a target of DCS, previous studies have shown that DCS competitively inhibits both Alanine racemase (Alr) and D-alanine-D-alanine ligase (Ddl) ^6,11^. However, more recent metabolomic study showed that Ddl is a primary target of DCS that is preferentially inhibited over alanine racemase (Alr) in *M. tuberculosis ^12^*. In addition, CycA is a transporter protein of D-alanine, D-serine and glycine D-serine/alanine/glycine ^13^, and its single nucleotide polymorphism (SNP) may partially contribute to the natural resistance to D-cycloserine in BCG ^10,14^. *ald* (*Rv2780*), encoding L-alanine dehydrogenase, was the fourth gene in *M. tuberculosis*, whose mutation was found in DCS resistant clinical isolates ^10^. However, mutations in *cycA* and *ald* only contribute to very low level resistance, and Ddl mutation was rarely found in studies without known phenotype in MDR/XDR-TB strains ^10^.

Despite the above progress and the significant advancements in general about the molecular understanding of drug resistance mechnanisms in *M. tuberculosis ^15^*, the distribution and characterization of DCS resistance in clinical strains are still vague. This is partly due to the technical difficulties with DCS susceptibility testing such that routine phenotypic assay is not performed in clinical labs ^16, 17^, as well as poor understanding of the molecular basis of resistance to this drug.

To better understanding the mechanisms of DCS resistance and to develop more rapid molecular tests for detection of its resistance, we characterized 35 DCS-resistant mutants isolated in vitro from *M. tuberculosis* H37Rv and discovered a panel of new unique mutations that are associated with DCS resistance that have not previously been reported.

## Materials and methods

### DCS resistant mutant isolation

One-month-old *M. tuberculosis* H37Rv cultures grown in 7H9 liquid medium supplemented with 0.05% Tween 80 and 10% bovine serum albumin-dextrose-catalase (ADC) enrichment were plated on 7H11 plates containing 20, 40, 80, 160, 320 mg/L cycloserine. After incubation at 37°C for 4 weeks, the mutant colonies were picked to confirm the drug resistance phenotype by transferring the mutants onto new plates containing different concentrations of DCS (20, 40, 80, 160 mg/L).

### Drug susceptibility testing

Drug susceptibility testing of the DCS-resistant mutants was performed on 7H11 agar plates containing 0, 20, 40, 80, 160 mg/L cycloserine *M. tuberculosis* H37Rv was included as a drug-susceptible control and BCG was included as a resistant control. The susceptible strain H37Rv did not grow on DCS-containing plates and only mutants that were consistently resistant to DCS were further analyzed by PCR and whole genome sequencing (WGS), as described below.

### Whole genome sequencing analysis

Genomic DNA from 18 cycloserine-resistant mutants, as well as the parent strain *M. tuberculosis* H37Rv, was sequenced using MiSeq (Illumina, Inc.) as described previously ^18^, except that paired-end sequencing libraries were constructed using Nextera XT DNA Sample Preparation kits (Illumina, USA) following manufacturer’s instruction. For each isolate, 500 M to 1.5 G bases (110-fold to 350-fold genome coverage) sequences were generated after barcodes were trimmed. Single-nucleotide variants (SNVs) and insertions and deletions (InDels) ranging from 1 to 5 bp were sorted and called at a minimum coverage of 4 reads using *M. tuberculosis* H37Rv genome (NC_018143.1) as a reference. Mutations in PE/PPE family genes and regions having repetition sequences were excluded from the analysis. Mutations in the parent strain *M. tuberculosis* H37Rv comparing with the genome online (NC_018143.1) were also excluded from the analysis.

### PCR and DNA sequencing

The genomic DNA from DCS-resistant mutants isolated in vitro was then subjected to PCR amplification using primers listed in Table 1. The PCR products were obtained using the amplification parameters listed in Table 1 and sequenced by Sanger method to confirm the mutations in these genes in selected mutants.

**Table 1.**
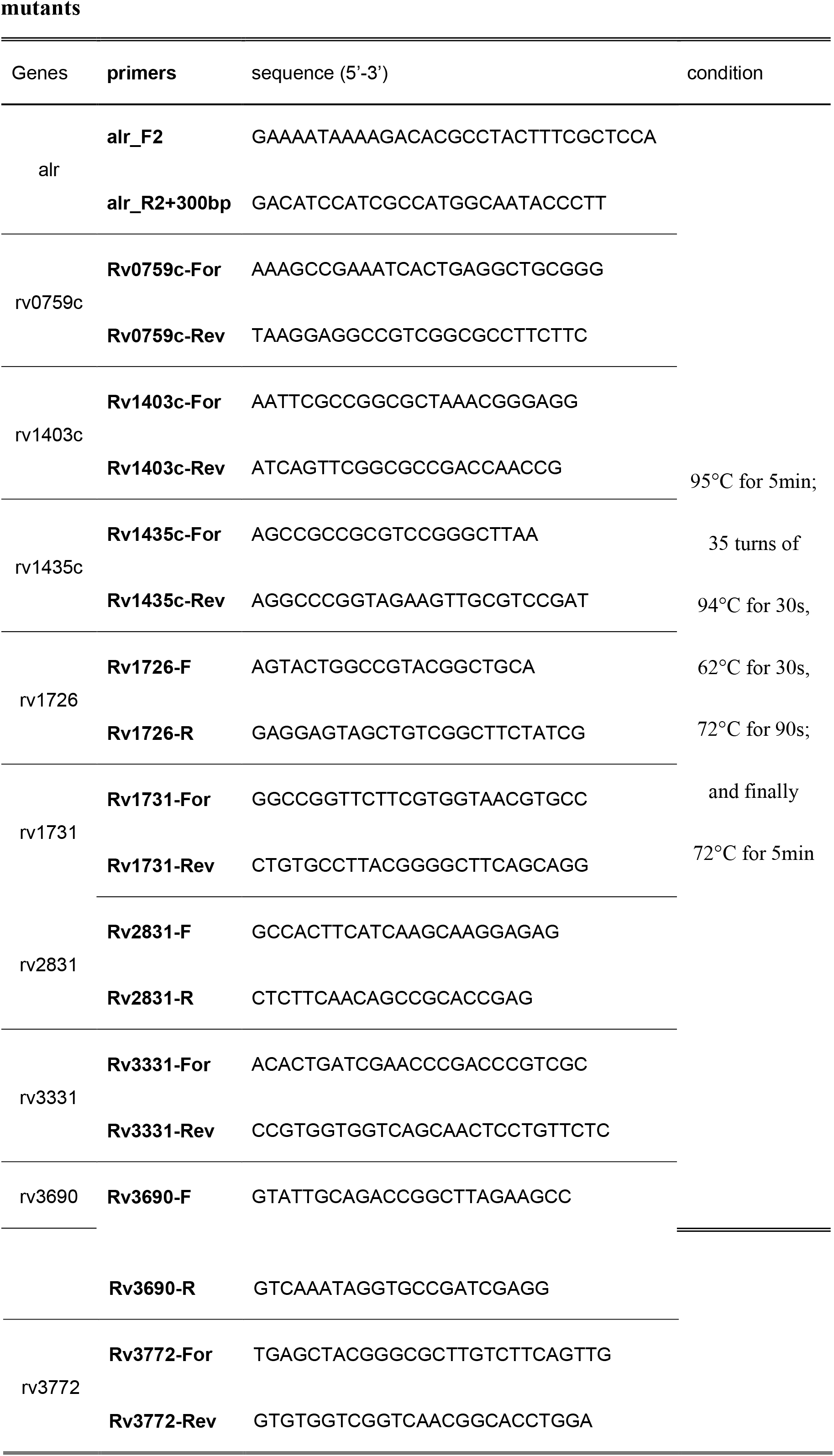
Primers and PCR conditions used to verify gene mutations in DCS resistant mutants

## Results and discussion

### Isolation of *M. tuberculosis* H37Rv mutants resistant to DCS

The cycloserine MIC for the sensitive *M. tuberculosis* H37Rv parent strain was found to be below 20 mg/L. To isolate mutants resistant to DCS, about 10^8^ *M. tuberculosis* bacteria were plated on 7H11 plates containing different concentrations of DCS (20, 40, 80, 160, 320 mg/L). After 4 weeks of incubation, no mutants grew on plates containing DCS higher than 80 mg/L. We found that only two mutants DT61-1 and DT69-1 grew on plates containing 40 mg/L cycloserine, and 35 mutants were obtained on plates containing 20 mg/L cycloserine. The mutation frequency of resistant mutants to 20 mg/L cycloserine was found to be about 2 x 10^−8^.

### Mutations identified in DCS resistant mutants by WGS

WGS of 18 DCS resistant mutants showed that 16 isolates had only 1 mutation (SNV or InDels) and 2 isolates had 2 mutations. Totally 17 different gene mutations were identified in 18 mutants (Table 2), and it is of interest to note that none of these mutations were dominant. Nonsynonymous mutations in *alr* (*rv3423c*)*, rv0059, betP* (*rv0917*)*, rv0221, rv1403c, rv1683, rv1726, gabD2* (*rv1731*)*, rv2749, sugI* (*rv3331*)*, hisC2* (*rv3 772*), single mutation in 5’ intergenic region of *rv3345c* and *rv1435c,* and insertion in 3’ region of *rv0759c* were identified as the solo mutation in their respective cycloserine-resistant mutants, suggesting these mutations associated with cycloserine resistance are highly diverse. In addition, one mutant (DT61-1) had two mutations in *alr (rv3423c)* as well as a −52 G-T change in *rv3345c* (hypothetical protein), while the other mutant (DT3-2) had double mutations in both *rv2831* and *rv3690* (Table 2). Ten mutation genes, *rv3690, rv0739c, rv1403c, rv1435c, rv2831, rv1726, rv1731, rv3331, rv3 772* and *alr* were verified by the Sanger sequencing method using primers and PCR conditions as described in Table 1.

**Table 2.**
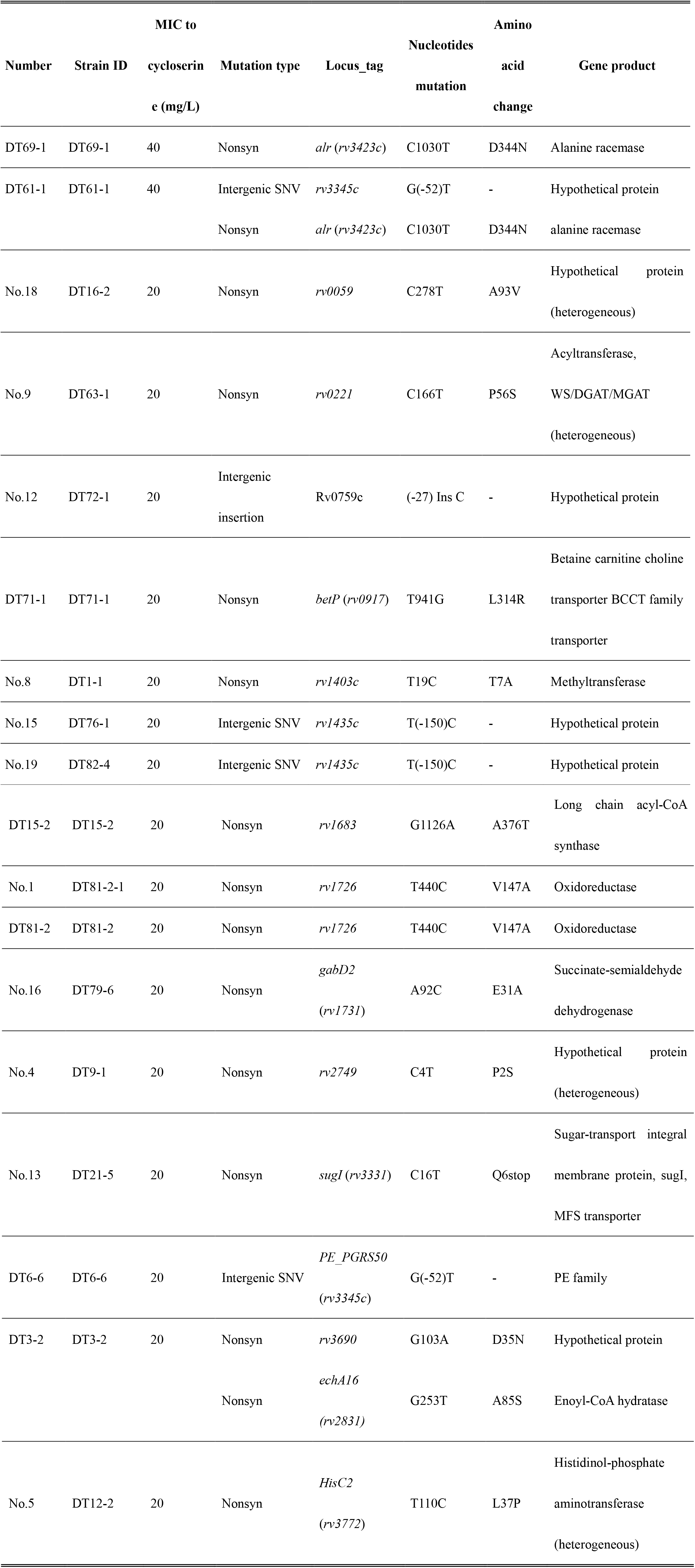
Mutations identified in cycloserine resistant mutants of *M. tuberculosis* by whole genome sequencing analysis

The two mutants DT61-1 and DT69-1 which had higher resistance to DCS (40 mg/L) had the same nonsynonymous mutation in alanine racemase gene *alr,* (nucleotide change C1030T, causing amino acid change of D344N). Except for *alr*, which is known to be involved in DCS resistance ^6,7^, the remaining 16 are novel and have not been reported previously. These 16 mutations include 3 lipid metabolism (Rv0221, Rv1683, Rv2831), 2 transport proteins (BetP, SugI), 1 toxin/antitoxin (Rv0059), 3 intermediary metabolism and respiration (Rv1726, gabD2, HisC2), 1 methyltransferase (Rv1403c), 1 PE family protein (PE_PGRS50), and 5 unknown hypothetical proteins (Rv0759c, Rv1435c, Rv2749, Rv3345c, Rv3690). *rv0059* is toxin of a TA-module (Rv0059 and Rv0060) ^19^.

One mutant (DT21-5) had an N-terminal stop codon mutation in the *sugI* gene, which resulted in complete loss of its protein function (Table 2). The *sugI* encodes a probable sugar-transport integral membrane protein in *M. tuberculosis* ^20^. Since the cycloserine structure is similar to natural furanose and *sugI* is the solo mutation detected in the genome, DCS could use SugI as the transporter for intake into the cell. The loss of function mutation in *sugI* could result in a lower uptake of cycloserine inside the cell and therefore leading to higher resistance to DCS. This is consistent with the previous observation that a transport protein involved in alanine and serine uptake was implicated in the uptake and resistance of DCS ^5^. *betP* was the other transport protein whose SNV was detected in a different mutant DT71-1. BetP transports molecules with a quaternary ammonium group like betaine, carnitine and choline. Whether mutations in SugI and BetP could cause resistance through alterating the transport of DCS remains further investigation in future studies.

Five genes encoding hypothetical proteins were identified in the mutants, including intergenic SNVs of *rv3345c, rv0759c* and *rv1435c,* and non-synonymous SNVs of *rv3690* and *rv2749.* Rv3345c seems to be a stress response protein, as it is regulated by *sigD* in strain H37Rv ^21^. The mutation is located in the promoter region of *rv3345c,* which may alter its expression. Rv3690 is a probable conserved membrane protein, and its expression was found to be elevated in MDR-TB strains ^22^. Although the role of *rv0759c* is not known, *rv0759c* may be an important gene in stress response. The intergenic region of this gene was predicted to be directly bound by SigF, SigC and SigK ^23^, and the control by SigF and SigH in heat stress were verified by Q-PCR ^24, 25^. Future studies are needed to address the role of the 5 intergenic regions in genes involved in stress response and membrane ptoteints in causing DCS resistance.

Rv1403c together with Rv1405c are important methyltransferase in *M. tuberculosis.* Their expression was very low under *in vitro* non-stressed and *ex vivo* conditions, but highly upregulated under *in vivo* and stressed conditions *in vitro* ^26^. They are both tightly regulated by the product of the *Rv1404* gene, encoding a member of the MarR family of transcriptional regulators and the stress condition upregulated their expression included acid shock, antibiotics like thioridazine, detergent like sodium dodecyl sulfate, hypoxic conditions and macrophage stress ^26–30^. Mutation in *rv1403c* impaired its growth phenotype in acidified 7H9 medium (pH5.5) compared with the wild type strain ^30^. *rv1726* encodes an FAD-binding dehydrogenase and its mutation was identified in 2 of 16 resistant mutants (DT81-2 and DT81-2-1) (Table 2). This oxidoreductase is structurally similar to 6-hydroxy-D-nicotine oxidase (6-HDNO) of *Arthrobacter oxidans* (29.5% identity), which oxidizes 6-hydroxy-D-nicotine to 6-hydroxy-N-methylmyosmine.

*rv3 772* encodes a histidinol-phosphate aminotransferase (HisC2), involved in the histidine-biosynthetic pathway. The pathway leads to enzymatic synthesis of histidine from 5-phosphoribosyl-1-pyrophosphate in ten steps, which is among the essential pathways required for optimal growth of *M. tuberculosis* and is conserved in archaea, bacteria, fungi and plants but not in mammals ^31,32^. The gene was shown to be involved in the adaptation to stress response and host cell environment, as it is regulated by FurA, an oxidative stress sensing regulator, and upregulated by rhodanine agent (D157070) in *M. avium* ssp. *paratuberculosis^33^’^34^*. *rv1731* encodes a succinate-semialdehyde dehydrogenase (GabD2), which catalyzes the NAD(P)^+^-coupled oxidation of succinic semialdehyde to succinate, the last step of the γ-aminobutyrate (GABA) shunt ^35^.

Three non-synonymous mutations in 3 lipid metabolism genes, including *rv0221* encoding a verified triacylglycerol synthase ^36^, *rv1683* encoding a long chain acyl-CoA synthase and *rv2831* encoding an enoyl-CoA hydratase were identified in 3 different mutants. *rv0221* belongs to the RD10 of *M. bovis* BCG and is located near LipC (*Rv0220*), lipW (*Rv0217c*), acyl-CoA synthetase (*Rv0214*), acyl-CoA dehydrogenase (*Rv0215c*), and an integral membrane acyltransferase (*Rv0228*). These genes may be cotranscribed under specific stimuli and may release fatty acid from triacylglycerol, carry out the transport of fatty acids and catalyze the resynthesis of triacylglycerols in the pathogen ^36,37^ *rv0221* was demonstrated up-regulated in intraphagosomal lesions and under multiple stresses ^38, 39^. The acyl-CoA synthase *Rv1683* is suspected to be essential for triacylglycerol hydrolysis and growth ^37, 40^, and was shown to be a functional esterase in all active, dormant and reactivation culture conditions ^41^. Rv2831 encodes an enoyl-CoA hydratase/isomerase family protein (EchA16), which is up-regulated in clinical MDR-TB strains and under exposure to mefloquine ^22,42^. As the triacylglycerol storage is important for the bacteria to transform into the dormancy-like state *in vitro* and survive during starvation, these three genes could be involved in bacterial dormancy and prolonged survival under stress. In this study, the mutations in these lipid metabolism genes may alter their enzyme activity, turn the bacteria into dormancy and therefore increase the survival against cycloserine.

Previous studies have reported only a few isolated DCS mutants for identifcation of molecular basis of DCS resistance ^5, 14^ This study analyzed the largest number of DCS resistant mutants so far. It is worth noting that among the known DCS resistance associated genes, we only found a mutation in *alr* in 2 of 18 DCS-resistant mutants (11%) but in none of the other known DCS associated resistance genes *ddl,* or *cycA* or *ald.* Instead, we found mutations in 16 novel genes that are associated with DCS resistance. Our finding of *alr* mutation in DCS resistant mutants is consistent with a previous study that demonstrated Alr being a primary target of DCS ^8^ but is in contrast to a recent study suggesting Ddl being a target of DCS ^12, 43^. However, this could be a reflection of the relatively small number of strains (18 mutants) being analyzed. Future investigations on more strains are required to better determine the frequency of the mutations in *alr, ddl,* as well as the new genes we identified in this work in DCS resistant clinical strains. In addition, the role of the 16 novel genes in DCS resistance should be addressed by molecular studies such as overexpression as well as point mutation constructions in future studies.

## Conclusions

In conclusion, we identified novel mutations associated with DCS resistance in *M. tuberculosis.* Our findings indicate that the mechanisms of DCS resistance are quite complex in *M. tuberculosis* and involve genes in lipid metabolism, methyltransferase, stress response, small transport proteins. This study provides useful information for improved understanding of molecular basis of DCS resistance and mechanisms of action recognition and also molecular detection of DCS resistance for improved treatment of MDR-TB. Future studies are needed to validate our findings in clinical isolates with DCS resistance and to address the role of the newly identified mutations in causing DCS resistance in *M. tuberculosis.*

## Transparency declarations

None to declare.

